# An allelic series at the *EDNRB2* locus controls diverse piebalding patterns in the domestic pigeon

**DOI:** 10.1101/2023.07.26.550625

**Authors:** Emily T. Maclary, Ryan Wauer, Bridget Phillips, Audrey Brown, Elena F. Boer, Atoosa M. Samani, Michael D. Shapiro

## Abstract

Variation in pigment patterns within and among vertebrate species reflects underlying changes in cell migration and function that can impact health, reproductive success, and survival. The domestic pigeon (*Columba livia*) is an exceptional model for understanding the genetic changes that give rise to diverse pigment patterns, as selective breeding has given rise to hundreds of breeds with extensive variation in plumage color and pattern. Here, we map the genetic architecture of a suite of pigmentation phenotypes known as piebalding. Piebalding is characterized by patches of pigmented and non-pigmented feathers, and these plumage patterns are often breed-specific and stable across generations. Using a combination of quantitative trait locus mapping in F_2_ laboratory crosses and genome-wide association analysis, we identify a locus associated with piebalding across many pigeon breeds. This shared locus harbors a candidate gene, *EDNRB2*, that is a known regulator of pigment cell migration, proliferation, and survival. We discover multiple distinct haplotypes at the *EDNRB2* locus in piebald pigeons, which include a mix of protein-coding, noncoding, and structural variants that are associated with depigmentation in specific plumage regions. These results identify a role for *EDNRB2* in pigment patterning in the domestic pigeon, and highlight how repeated selection at a single locus can generate a diverse array of stable and heritable pigment patterns.

**AUTHOR SUMMARY:** Both wild and domestic birds show striking variation in pigment patterning, and these pigment patterns can play critical roles in mate choice, communication, and camouflage. Despite the importance of pigment patterning for survival and reproductive success, the mechanisms that control pigment patterning remain incompletely understood. In domestic pigeons, artificial selection has given rise to a wide array of pigmentation patterns within a single species, including a suite of phenotypes called “piebalding” that is characterized by regional loss of feather pigment. Here, we took advantage of the wide array of distinct piebalding phenotypes in domestic pigeons to map the genetic basis of region-specific loss of plumage pigment. Rather than focusing on a single piebalding pattern, we sought to broadly understand genetic control of regional pigment loss by examining several breeds with different piebalding patterns. We compared the genomes of piebald and non-piebald pigeons using both genetic crosses and genome-wide association studies, and identified several genetic changes affecting the endothelin receptor gene *EDNRB2*. Our findings highlight how independent mutations at a single locus can drive diversification of plumage pigment patterning within a species.

## INTRODUCTION

Vertebrates display striking variation in both pigmentation types and patterning. In a natural setting, intraspecific pigment variation is often associated with sexual dimorphism or adaptation to different environmental conditions across a species range. Similarly, selective breeding in domestic species has given rise to a diverse array of pigment patterns [1,2]. Variation in pigment patterns within and among vertebrates reflects underlying changes in cell migration and function that can have profound impacts on health, behavior, and survival [2–5]. Thus, identifying molecular changes that underlie variation in pigment patterning is crucial to understanding both the evolutionary origins of diversity and the etiology of pigment-related developmental disorders.

One broad category of pigment patterning that has evolved repeatedly within and among species is piebalding. Piebald patterns are characterized by patches of pigmented and non-pigmented skin, hair, feathers, or scales. Among mammals, piebalding is linked to multiple pigmentation pathway genes, including *KIT* in horses and pigs*, EDNRB* in horses and mice, and *MITF* in dogs and cattle [6–12]. In addition to altering pigment patterning, genetic variants sometimes have broader pleiotropic effects. For example, disruption of *EDNRB* is associated with enteric nervous system defects, while changes at the *MITF* locus are linked to deafness [5,9,13,14]. Therefore, understanding the genetic and developmental mechanisms that control depigmentation phenotypes has critical relevance from developmental, evolutionary, and clinical perspectives.

Domestic pigeons (*Columba livia*) are an ideal model to study the genetics of plumage depigmentation and piebald patterning. For centuries, hobbyists have selected for a wide range of depigmentation phenotypes, ranging from “recessive white” birds that exhibit total loss of both plumage and iris pigmentation to an array of “piebald” markings, defined as any patchwork of white and colored plumage [15,16]. Some breeds, such as the Helmet, have mostly white plumage with patches of pigmented feathers restricted to the head and tail (Fig. 1A). Conversely, other breeds like the Mookee have very little white plumage, limited to the head and the primary flight feathers (Fig. 1B). The inheritance patterns of piebalding also vary. Crosses between piebald and non-piebald breeds show that some piebald patterns have dominant inheritance, others are recessive, and some show variability in plumage patterning among F_1_ offspring [15]. Based on analyses of phenotypes from a variety of crosses, prior work implies that different genetic factors control piebald patterning in different body regions, and that some combinations of depigmented plumage are caused by multiple linked variants on the same chromosome [15,17].

**Figure 1.**
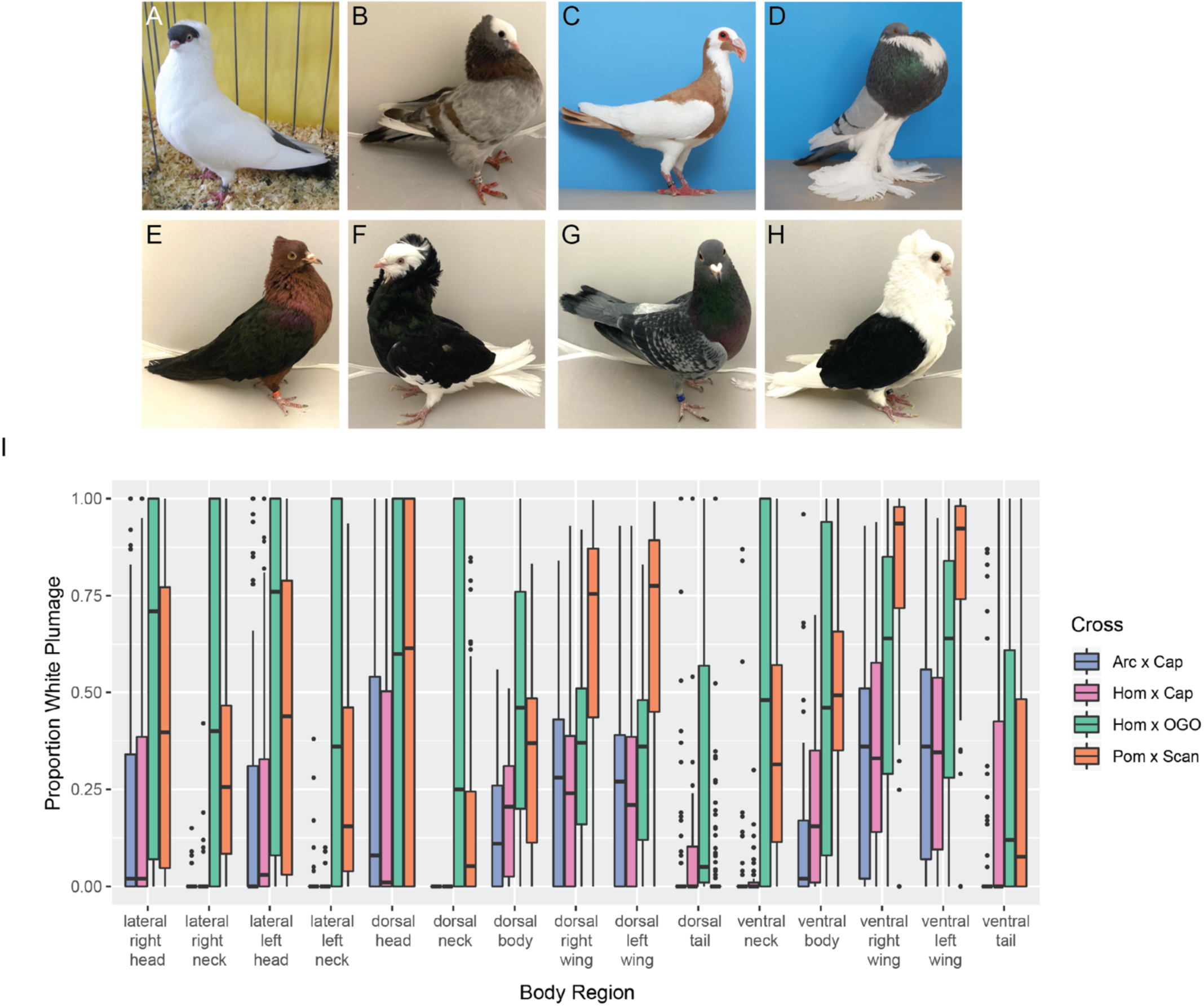
Piebald patterns vary among breeds and within F_2_ crosses. (A-H) Examples of piebald and cross founder breeds. (A) Helmet, (B) Mookee, (C) Scandaroon (Pom x Scan founder breed), (D) Pomeranian Pouter (Pom x Scan founder breed), (E) Archangel (Arc x Cap founder breed), (F) Old Dutch Capuchine (Arc x Cap and Hom x Cap founder breed), (G) Racing Homer (Hom x Cap and Hom x OGO founder breed), (H) Old German Owl (Hom x OGO founder breed). E and F are images of founders of the Arc x Cap cross, others are representative images. (I) Boxplots showing the distribution of the proportion of white plumage for F_2_ birds from each cross in the 15 different body regions quantified. Boxes span from the first to third quartile of each data set, with lines indicating the median. Whiskers span up to 1.5x the interquartile range. Photos in (C) and (D) by Layne Gardner, used with permission.

In this study, we use parallel genetic, genomic, and developmental biology approaches to test these genetic. We previously used quantitative trait locus (QTL) mapping in an F_2_ intercross to identify two loci that contribute to piebalding [18]. Here, we evaluate three additional crosses to identify similarities and differences in both phenotypes and the genetic architecture of piebalding in additional breeds. Next, we examine whole genome sequences from a wide array of pigeon breeds to test for associations between genotypes and piebalding phenotypes. Through these complementary approaches, we identify both breed-specific and shared loci associated with piebalding. At a major locus associated with piebalding in multiple breeds, we identify a candidate gene, *EDNRB2*, with a mix of coding, noncoding, and structural variants. Our combined genetic, genomic, and developmental results indicate that multiple variants at the same locus likely produce the diverse suite of phenotypes that breeders have selected under domestication.

## RESULTS

### Piebald phenotypes vary among laboratory crosses

As a first step in mapping the genetic architecture of piebalding, we examined phenotypes in three F_2_ crosses, each with one piebald and one non-piebald founder. The first cross was derived from a non-piebald Archangel and a piebald Old Dutch Capuchine (Arc x Cap), the second from a non-piebald Racing Homer and a piebald Old Dutch Capuchine (Hom x Cap), and the third from a non-piebald Racing Homer and a piebald Old German Owl (Hom x OGO). In addition, we compared these three crosses to a previously analyzed a cross derived from two piebald founder breeds, the Pomeranian Pouter and the Scandaroon (Pom x Scan) [18]. Comparing piebalding in crosses with diverse piebald founders allows us to identify potential differences in the genetic basis of this trait among breeds.

We quantified the proportion of white and pigmented plumage across 15 different body regions in each cross (Fig. 1I). For most body regions, we observed overlap in the range of plumage phenotypes observed among crosses; however, we found differences in both the extent and distribution of white plumage in some body regions. Much of this variation is consistent with expectations based on founder phenotypes. For example, in the Arc x Cap and Hom x Cap crosses, we never observed white plumage on the dorsal neck, but fully white dorsal necks are common in the Hom x OGO cross. The Old German Owl founder of the Hom x OGO cross has a fully white neck, while the Old Dutch Capuchines have pigmented dorsal neck plumage and limited white feathering on the ventral and lateral neck. In another example, the extent of white feathering on the wings is greater in the Pom x Scan cross compared to other crosses. Consistent with the wing phenotype of F_2_s, the Scandaroon founder breed has a broader region of white plumage on the dorsal wing compared to the Old German Owl or Old Dutch Capuchine (see Fig. 1 C-H).

Principal component analysis (PCA) of piebalding phenotypes shows overlap in the phenotypic space occupied by all four crosses. One exception is a subset of Hom x OGO F_2_ offspring that separate from other pigeons on both PC1 and PC2, which reflect the level of white plumage in different body regions (Supplemental Fig. S1, blue triangles at lower right). These individuals have more extensive white plumage than the most extreme phenotypes in other crosses, with many closely recapitulating the Old German Owl founder breed (Fig. 1H). While variability both within and among crosses aligns well with expectations based on founder phenotypes, the founder phenotypes do not strictly define the limits of piebalding in F_2_ birds. Several F_2_ individuals have white plumage in body regions where cross founders do not (Supplemental Fig. S2A-B), suggesting that either the effects of genetic modifiers not visible in the phenotypes of the founders, stochasticity, or both may play a role in determining the extent of piebalding.

We next sought to determine if piebalding consistently co-occurred in two or more specific body regions in our crosses (Supplemental Fig. S2), which might suggest shared genetic control of those regions. We found that in the Arc x Cap and Hom x Cap crosses, the strongest correlations are within body regions (e.g., lateral head and dorsal head, dorsal left wing and dorsal right wing), with the head, wing, and tail forming independent “modules” that separate out over PC2. In the Hom x OGO cross, correlation is high across most body regions, and PC2 is primarily defined by tail pigmentation. In the Pom x Scan cross, we identified two sets of correlated regions, one comprising the neck and body and the other comprising the head and the wing. This result is similar to our previous findings for this cross that identified two major QTLs for piebalding, one contributing primarily to white plumage on the ventral neck and dorsal body, and the other contributing to white plumage on the wings and head [18].

In summary, we found inconsistent linkages between piebald body regions among the F_2_ progeny of four lab crosses. These patterns could arise from multiple piebalding-associated genetic variants, some of which affect plumage depigmentation in a single body region and some of which drive depigmentation across multiple body regions.

### Shared and cross-specific loci control plumage depigmentation

We next asked which regions of the genome affected plumage depigmentation. We used QTL mapping to identify loci associated with the proportion of white plumage across 15 different body regions in the Arc x Cap, Hom x Cap, and Hom x OGO crosses (Fig. 2, Figs. S3-S5, Table 1). A major-effect QTL on LG15 was associated with piebalding across many body regions in all three new crosses and this locus was also a major-effect QTL in the Pom x Scan cross [18].Thus, a single locus drives a substantial proportion of the variation in piebald patterning in multiple breeds (Fig. 3, Table 1). The boundaries of QTL intervals vary both among crosses and body regions: the union of 2-LOD intervals from the peak marker for all crosses and body regions spans 14.7 Mb and 11 genomic scaffolds of the Cliv_2.1 assembly. However, the intersection of all 2-LOD intervals in all crosses spans only 162 kb on scaffold 507 (NCBI accession AKCR02000023.1) and contains two annotated genes — *RPAC2* and *EDNRB2* — and 128 kb of intergenic space 5’ of *EDNRB2*.

**Figure 2.**
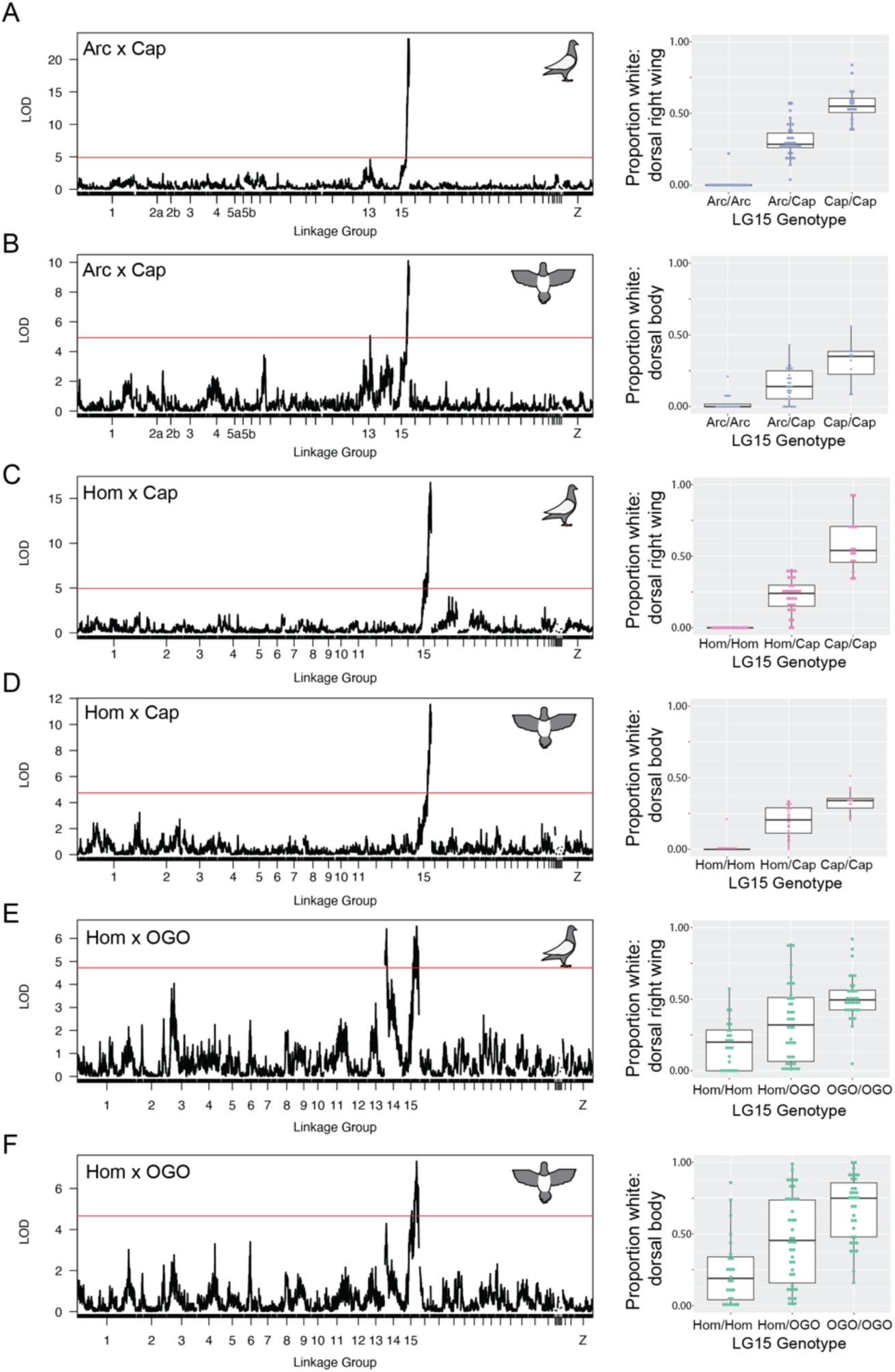
QTL mapping identifies shared and cross-specific regions associated with piebalding. Representative QTL results for piebalding in the Arc x Cap (A-B), Hom x Cap (C-D), and Hom x OGO (E-F) crosses. A,C, and E show QTL results for the dorsal right wing. B, D, and F show QTL results for the dorsal body. Lefthand plots illustrate QTL mapping results, with linkage map position on the X-axis and LOD score on the Y-axis. Red lines indicate the threshold for genome-wide statistical significance. Righthand boxplots illustrate the relationship between genotype at the LG15 peak marker and the proportion of white plumage in that body region. Boxes span from the first to third quartile of each data set, with lines indicating the median. Whiskers span up to 1.5x the interquartile range. QTL mapping results for additional body regions are in Supplemental Figures S2, S3, and S4.

**Figure 3.**
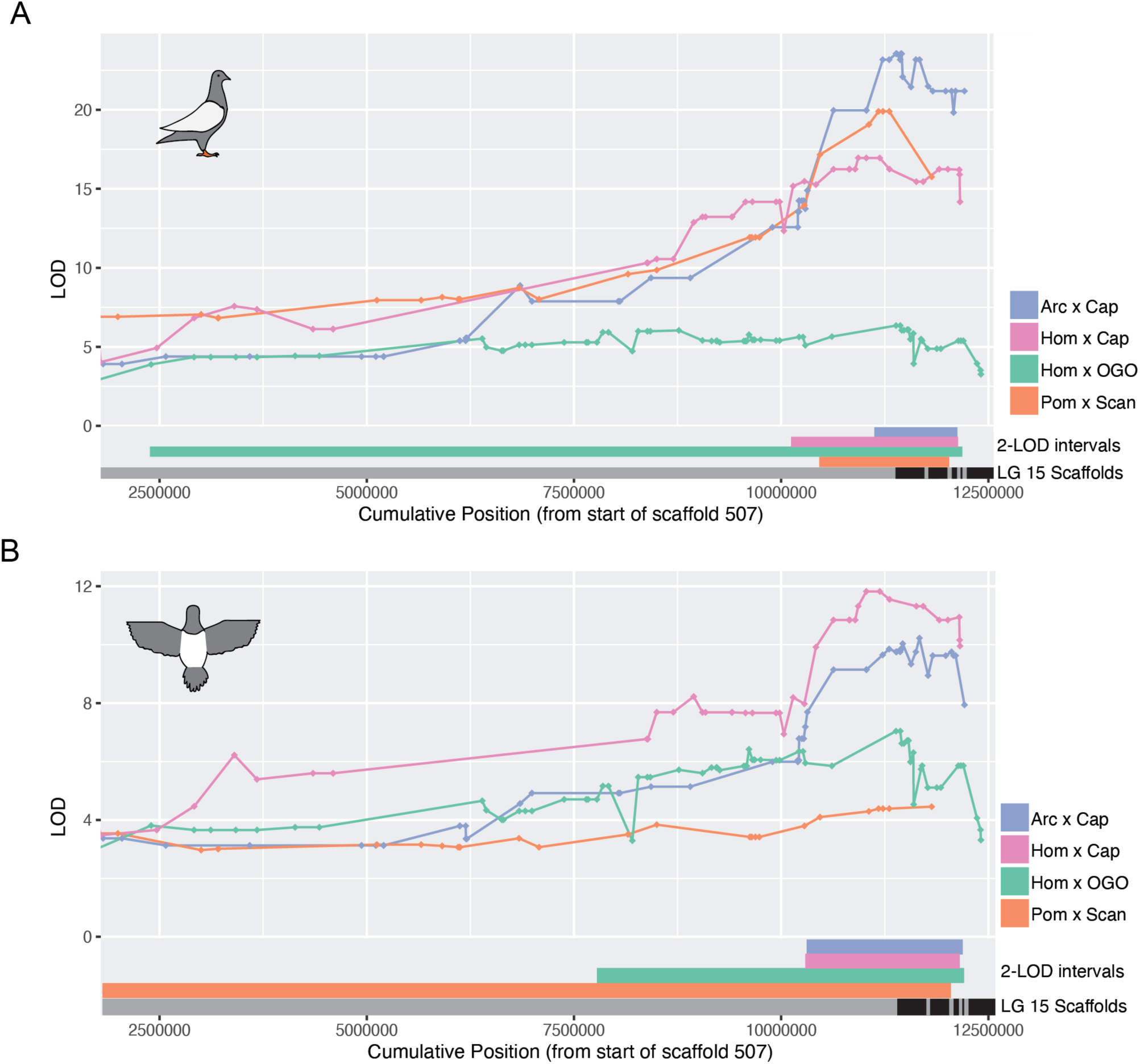
Piebalding QTLs on linkage group 15 overlap in all crosses. Overlaid QTL results for dorsal right wing (A) and dorsal body (B) within the LG15 peak region. 2-LOD intervals for each cross are indicated at the bottom of each plot. Black and gray bars indicate scaffold boundaries for the scaffolds that make up the candidate region in all four crosses; some scaffolds are not represented in all crosses due to lack of informative markers.

**Table 1.** Summary of QTL analysis for regional white plumage in the Arc x Cap, Hom x Cap, and Hom x OGO crosses. PVE, percent variance explained.

|  | Arc x Cap |  |  |  | Hom x Cap |  |  |  | Hom x OGO |  |  |  |
| --- | --- | --- | --- | --- | --- | --- | --- | --- | --- | --- | --- | --- |
| Body Region | Linkage Group | Peak Marker | Peak LOD score | PVE | Linkage Group | Peak Marker | Peak LOD score | PVE | Linkage Group | Peak Marker | Peak LOD score | PVE |
| Lateral Right Head | 15 | 1916:103567 | 18.5 | 70.9 | 15 | 507:9653061 | 20.9 | 81.1 | 15 | 1916:291786 | 6.5 | 25.7 |
| Lateral Right Neck | Not significant |  |  |  | Not significant |  |  |  | 14 | 1259: 880682 | 4.9 | 16.1 |
|  |  |  |  |  |  |  |  |  | 15 | 507:11385801 | 6.6 | 21.8 |
| Lateral Left Head | 15 | 683.1: 222122 | 15.4 | 64.1 | 15 | 507:10820150 | 25.2 | 86.5 | 15 | 1916:291786 | 6.5 | 25.7 |
| Lateral Left Neck | Not significant |  |  |  | Not significant |  |  |  | 14 | 1259: 880682 | 5.9 | 18.9 |
|  |  |  |  |  |  |  |  |  | 15 | 507: 11385801 | 6.3 | 20.3 |
| Dorsal Head | 15 | 1916:147181 | 13.8 | 60.2 | 15 | 507:10820150 | 17.3 | 74.7 | 15 | 1916: 291786 | 5.1 | 20.8 |
| Dorsal Neck | No depigmentation in this region |  |  |  | No depigmentation in this region |  |  |  | 14 | 1259: 880682 | 5.8 | 18.5 |
|  |  |  |  |  |  |  |  |  | 15 | 1916:291786 | 6.2 | 19.7 |
| Dorsal Body | 13 | 1486: 1732159 | 5.1 | 9.1 | 15 | 507:11027392 | 11.5 | 60.0 | 15 | 507:11385801 | 7.3 | 28.4 |
|  | 15 | 1916:103567 | 10.1 | 29.4 |  |  |  |  |  |  |  |  |
| Dorsal Right Wing | 15 | 507:11385801 | 23.2 | 79.7 | 15 | 507:11027392 | 16.7 | 73.6 | 14 | 1259: 1259791 | 6.4 | 26.0 |
|  |  |  |  |  |  |  |  |  | 15 | 507:11385801 | 6.5 | 26.4 |
| Dorsal Left Wing | 13 | 1486: 1188321 | 5.48 | 1.1 | 15 | 507:10912510 | 15.3 | 70.3 | 14 | 1259: 844674 | 5.5 | 19.7 |
|  | 15 | 507:11385801 | 24.2 | 50.6 |  |  |  |  | 15 | 507:11385801 | 8.1 | 28.4 |
| Dorsal Tail | Not significant |  |  |  | Not significant |  |  |  | 14 | 1259: 1259791 | 8.2 | 27.1 |
|  |  |  |  |  |  |  |  |  | 15 | 507:9260009 | 4.8 | 15.8 |
| Ventral Neck | 13 | 493: 4593712 | 6.6 | 50.5 | 15 | 683.1: 144415 | 5.9 | 37.9 | 14 | 1259: 880682 | 5.9 | 19.5 |
|  | 15 | 507:10316297 | 5.0 | 43.5 |  |  |  |  | 15 | 507:11385801 | 5.7 | 18.7 |
| Ventral Body | 15 | 683.1_222122 | 8.7 | 43.9 | 15 | 507:11027392 | 15.3 | 70.4 | 14 | 1259_747998 | 4.9 | 16.6 |
|  |  |  |  |  |  |  |  |  | 15 | 507:11385801 | 7.4 | 25.3 |
| Ventral Right Wing | 15 | 1916:103567 | 23.9 | 79.8 | 15 | 507:11027392 | 14.1 | 67.3 | 15 | 507:10611804 | 7.5 | 29.1 |
| Ventral Left Wing | 13 | 1486: 1188321 | 5.2 | 1.2 | 15 | 507:11027392 | 12.9 | 64.2 | 15 | 507:11385801 | 7.9 | 30.3 |
|  | 15 | 507:11385801 | 25.6 | 53.9 |  |  |  |  |  |  |  |  |
| Ventral Tail | Not significant |  |  |  | 15 | 507:9409137 | 6.0 | 38.2 | 14 | 1259: 880682 | 8.6 | 27.3 |
|  |  |  |  |  |  |  |  |  | 15 | 507:9260009 | 4.9 | 14.9 |

While the LG15 QTL is shared between crosses, the Arc x Cap and Hom x OGO crosses also have QTLs associated with piebalding on other linkage groups that are cross-specific, indicating that piebalding phenotypes can be polygenic (Fig. 2B,E; Supplemental Fig. S3,S5). The Hom x OGO cross has a second major-effect QTL on LG14 that contributes to piebalding across many body regions (Fig. 2E, Supplemental Fig. S5), while the Arc x Cap cross has a minor effect QTL on LG13 that primarily impacts the ventral neck and dorsal body (Fig. 2B, Supplemental Fig. S3).

### A single genomic locus is consistently associated with piebalding in a wide variety of pigeon breeds

QTL mapping showed that piebalding is controlled by LG15 in several breeds; however, this method could not tell us whether these and other piebald breeds share the same genetic variants in the candidate region. Therefore, we examined allele frequency differentiation between the genomes of piebald (43 genomes, 31 breeds) and non-piebald (98 genomes, 44 breeds and feral pigeons) pigeons using probabilistic F_ST_ (pF_ST_) [19,20]. We identified a single well-differentiated peak on scaffold 507 that is strongly associated with piebalding (Fig. 4A). Overall, 296 SNPs on 14 scaffolds reached the threshold for genome-wide statistical significance. Of these, 265 (90%) are within a 373-kb region spanning scaffold 507:11066907-11440529, and 43 SNPs define a smaller 215-kb region that reaches the maximum pF_ST_ statistic.

**Figure 4.**
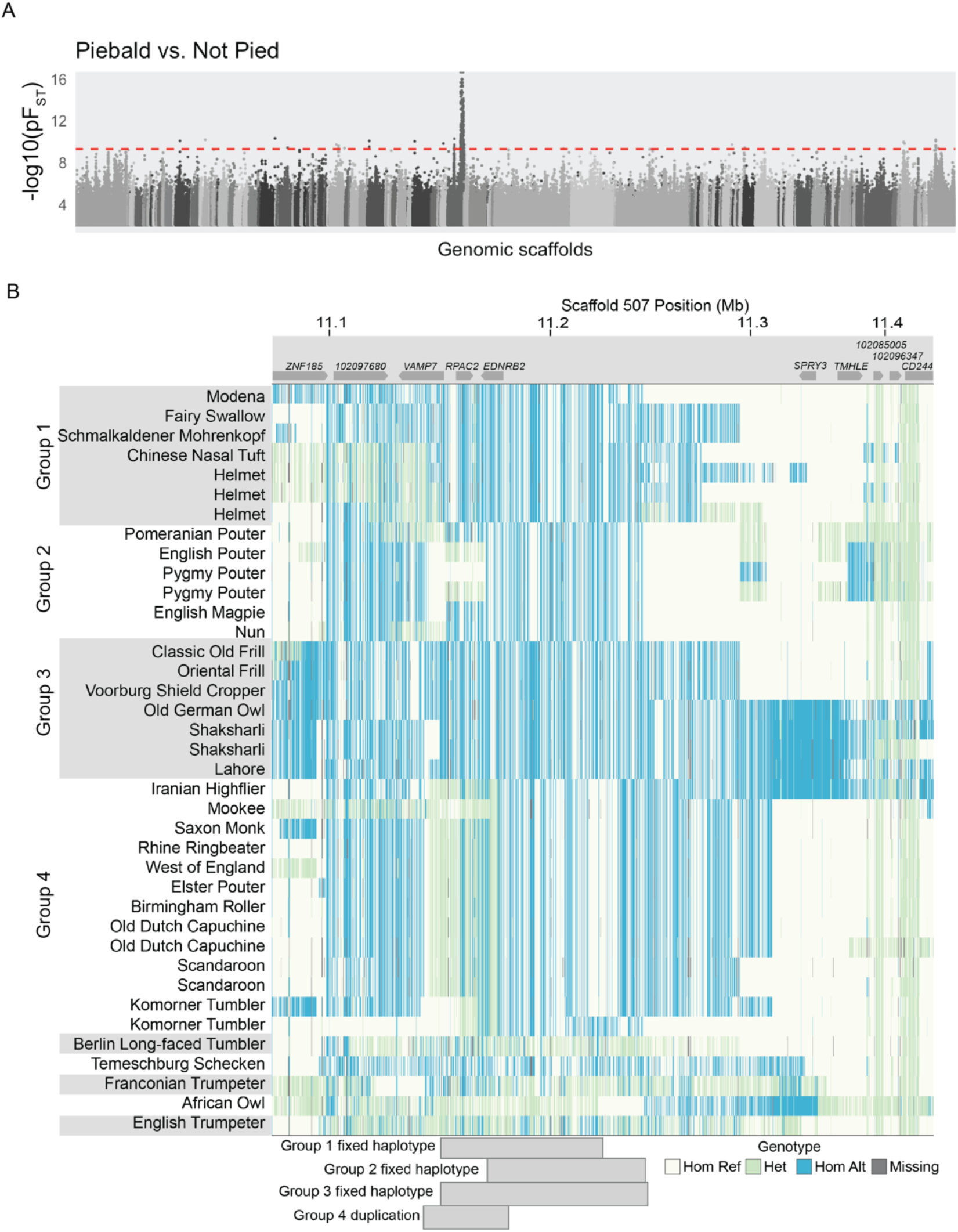
A single genomic locus associated with piebalding across many breeds. (A) Whole-genome pF_ST_ comparisons of piebald birds to non-piebald birds. Dashed red line indicates 5% threshold for genome-wide significance. (B) Plot of genotypes within the candidate region identified by pF_ST_. Each row shows an individual bird, and each vertical line a SNP position colored by genotype relative to the reference genome. Approximate locations of genes are illustrated at top. Shading on the left-hand side indicates birds with shared fixed haplotypes, the approximate span of these haplotypes is illustrated by gray blocks on the bottom of the plot. Hom Ref, homozygous reference allele; Het, heterozygous; Hom Alt, homozygous alternate (non-reference) allele.

The 215-kb region spans three complete genes (*VAMP7, RPAC2,* and *EDNRB2*), a portion of a fourth gene (*LOC102097680*, SLAIN motif-containing protein-like), and approximately 140 kb of intergenic space 5’ of *EDNRB2*. Therefore, this pF_ST_ candidate region converges on the same region identified by QTL mapping: it lies completely within the broadest interval defined by all four crosses and spans the entirety of the 162-kb minimal candidate QTL interval. Given the association of this region with piebalding in all four crosses and a diverse panel of genomes, we chose to focus on the shared LG15/scaffold 507 candidate interval as the major piebalding candidate region.

### Piebald pigeons do not share a single haplotype at the candidate locus

The candidate region on scaffold 507 is associated with piebalding in many breeds. However, the variance in phenotypes and inheritance among crosses raises the possibility of allelic heterogeneity at this locus. We examined sequences in the candidate region and found that, for many of the significant SNPs identified by pF_ST_, piebald-associated alleles are not shared across all piebald genomes. Instead, we identified four haplotype groups (Groups 1-4, below) with distinct patterns of shared SNP genotypes that correlate with piebalding in specific body regions (Fig. 4B, Supplemental Fig. S6).

Group 1 includes pigeons with white necks and bodies, partially-to fully-pigmented heads, and pigmented tails. Wing plumage in this group is variable: all birds have white flight feathers, but wing shield feathers can be pigmented or white (e.g., Fig. 1A). Group 2 consists primarily of pouter breeds characterized by pigmented heads, white ventral bodies and wing flights, and a U-shaped “bib” marking across the lateral and ventral neck (e.g., Fig. 1D). Group 3 contains a combination of piebald patterns characterized by white neck and breast plumage and variable piebalding on the head, ranging from patches of white feathers to fully white heads (e.g., Fig. 1H). Group 4 consists primarily of birds with patterns that breeders have termed “baldhead” or “monked” (e.g., Fig. 1B, F). All birds in this group are characterized by primarily white heads and white wing flights. Some also have white wing shields, but the scapular feathers, where the wings articulate with the shoulder, are pigmented (e.g., Fig. 1C). Below, we describe the unique genetic signatures at the candidate locus that define each group.

### Groups 1 and 2: coding mutations in *EDNRB2* are associated with plumage depigmentation

To assess group-specific allele frequency differentiation, we compared allele frequencies across the scaffold 507 candidate region in each subgroup to non-piebald birds by pF_ST_. For each group, we first looked for significantly differentiated SNPs, then identified piebald candidate SNPs within the differentiated region. The reference assembly is from a non-piebald bird; therefore, we defined piebald candidate SNPs as variants where the nonreference allele was fixed in the piebald group, but never present in the homozygous state in the non-piebald background genomes.

### Group 1: an R290C substitution is associated with piebalding

In Group 1 birds, we identified 31 piebald candidate SNPs spanning a 92 kb (scaffold 507: 11137571-11229275) (Fig. 5A). A non-reference variant at scaffold 507:11166924 is in exon 5 of *EDNRB2* and, as the only candidate SNP that is located with the coding region, is predicted to cause an amino acid substitution from arginine to cysteine at position 290 (R290C) in the EDNRB2 protein. EDNRB2 is a G-protein coupled receptor with seven transmembrane domains. The R290C substitution lies on the edge of the fifth transmembrane domain and within a putative peptide ligand binding pocket, as predicted based on the structures of ligand-bound family members (accession #cd15977) [21,22]. Protein alignments of the EDNRB2 sequence across five tetrapod species show that the R290 residue is not highly conserved, but this mutation is classified as “probably damaging” by PolyPhen2 (score=0.981; Fig. 5B) [23]. We next screened for this mutation in 47 additional pigeon genomes that were not included in the pF_ST_ analysis and identified one additional bird homozygous for the R290C mutation, an American Highflier. Based on the breed standard, this bird is expected to have a piebalding pattern similar to other birds with the R290C mutation [24].

**Figure 5.**
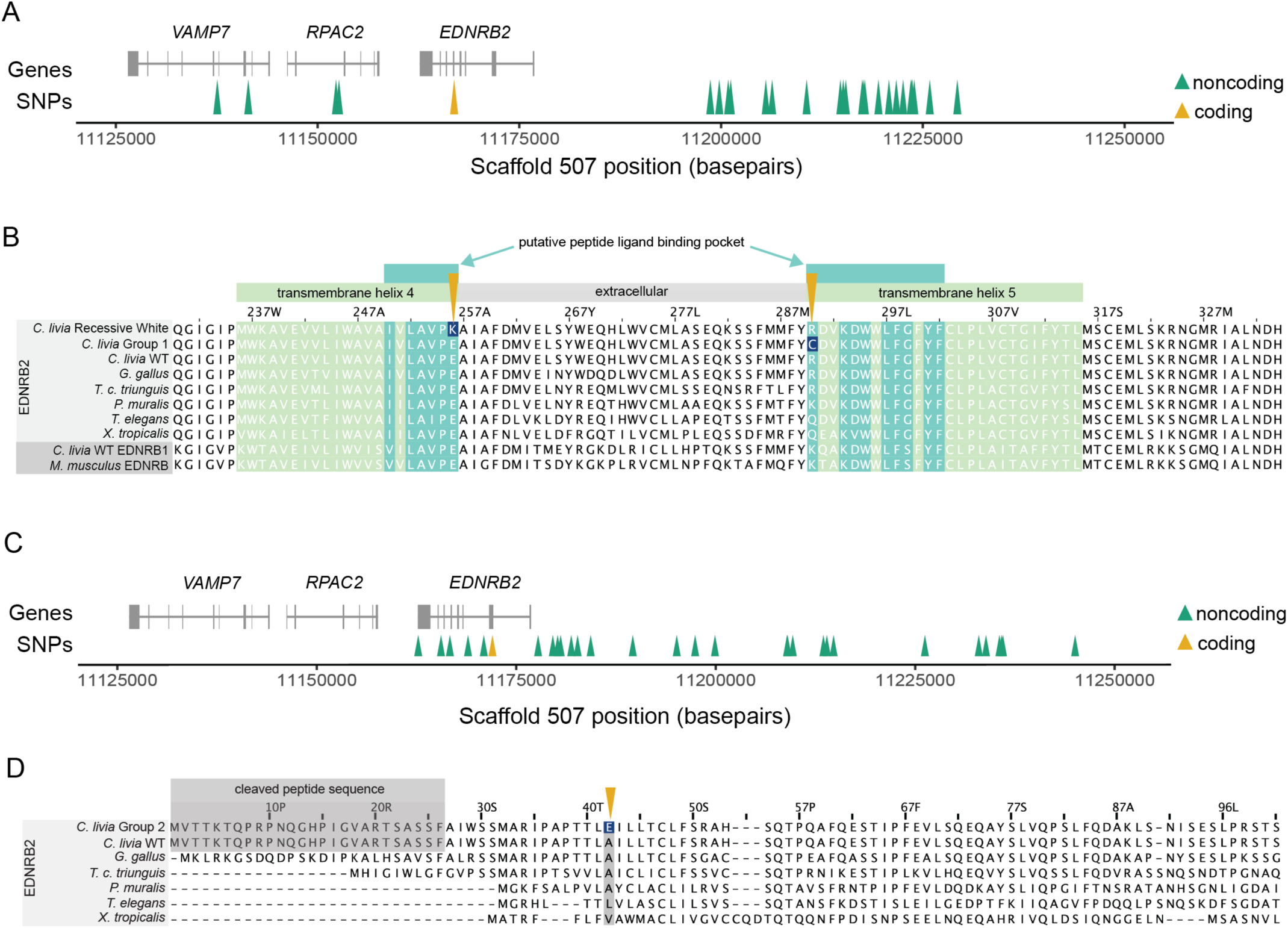
Three coding mutations are associated with plumage depigmentation. (A) Genomic context of the Group 1 piebald haplotype. Gene models for the region are shown in gray. Arrowheads indicate coding (yellow) and non-coding (green) SNPs. (B) Multi-species alignment of a portion of EDNRB2 (light gray rows) and EDNRB/EDNRB1 (dark gray, bottom two rows) protein sequences. Amino acids within a transmembrane helix are shaded in light green, residues that are part of a putative ligand binding pocket are shaded dark green. The coding changes identified in recessive white and Group 1 piebald birds, like the Helmet breed, are marked by yellow arrows, with the altered residues shaded dark blue. (C) Genomic context of the Group 2 piebald haplotype. (D) Multi-species alignment of a portion of EDNRB2 protein sequences. The cleaved peptide at the 5’ end of the *C. livia* protein is marked in gray; this portion exists in other species but is not easily delineated in this alignment due to variability of the amino end of the protein. The coding change identified in Group 2 birds is marked a yellow arrow, with the altered residue shaded dark blue.

### A E256K mutation in EDNRB2 is linked to recessive white

Our analysis of Group 1 pigeons unexpectedly led to the discovery of a coding mutation associated with a classic pigmentation trait known as “recessive white,” which is characterized by completely white plumage and “bull” (dark brown to black) eye color [15]. One of the Group 1 pigeons carrying the R290C mutation belongs to the Modena breed and has piebald markings known as “Gazzi”. Extensive crosses by pigeon breeders suggest that Gazzi (*z*) is allelic and dominant to recessive white (*z^wh^*) [15], suggesting that a *z^wh^* allele should also be present on a different haplotype at this locus. We examined the *EDNRB2* coding sequence of three bull-eyed white individuals from three different breeds and found a coding mutation in exon 4 (C to T at scaffold 507:11167700). Two of the three birds are homozygous and the third is heterozygous for this nucleotide change, which results in a glutamic acid to lysine substitution in at position 256 (E256K) of the EDNRB2 protein. Like the R290C mutation identified in Group 1 piebald birds, the E256K substitution is predicted to be “probably damaging” to the peptide ligand binding pocket (PolyPhen2 score=0.999) [23] (Fig. 5B). Unlike R290, however, E256 is highly conserved across vertebrates. In addition to its conservation in vertebrate EDNRB2 proteins, this residue is conserved in *C. livia* EDNRB1 and *M. musculus* EDNRB, suggesting it may be critical for the function of endothelin receptors broadly.

To increase our sample size, we used PCR to test for the candidate *z^wh^* mutation in 9 additional bull-eyed white pigeons from 7 different breeds. Of these, 6 were homozygous for the E256K mutation, 2 were heterozygous, and 1 did not carry the mutation. These results suggest that the E256K mutation is associated with bull eye color and white plumage, but it is not the only genetic variant that can cause the bull-eyed white phenotype. In bull-eyed white birds heterozygous for the E256K mutation, genetic background or the presence of other piebalding alleles may contribute to the phenotype. Indeed, through crosses, pigeon breeders have determined that while the recessive white phenotype is common, bull eye color and fully white plumage can also arise from other combinations of alleles, including combinations of multiple piebalding alleles [15].

### Group 2: an A42E substitution in EDNRB2 is associated with piebalding

In Group 2 pigeons, we identified 33 piebald candidate SNPs spanning 82 kb (scaffold 507: 11162719-11245041) (Fig. 4B, Fig. 5C). Only one of these Group 2 variants is a coding variant, which results in an alanine to glutamic acid substitution at position 42 (A42E) of the EDNRB2 protein (Fig. 5C). This region is predicted to be an extracellular domain between the signal peptide sequence and the first transmembrane domain, and multi-species alignments show partial conservation of this residue (Fig. 5D). This mutation is classified as “possibly damaging” (PolyPhen2 score = 0.86) [23].

In summary, we identified three coding mutations associated with piebalding in *EDNRB2.* Two are associated with the location and extent of piebalding in Group 1 and 2 pigeons that retain some pigmented plumage, and the third is associated with total loss of plumage pigment in recessive white birds. Two of these coding mutations, E256K and R290C, are in the same ligand binding pocket of the EDNRB2 protein but have markedly different effects on plumage phenotype. Based on amino acid characteristics, conservation of the mutated residues, and the phenotypes of birds harboring these mutations, we hypothesize that the E256K mutation in recessive white birds may create a null allele that disrupts pigment cell precursors throughout the embryo in the homozygous state. In contrast, the R290C mutation in Group 1 piebald birds may create a hypomorphic allele of EDNRB2 with altered binding affinity, leading to partial loss of pigment. These pigeons do not show total loss of plumage pigment, indicating that some level of function is retained. The A42E mutation identified in Group 2 birds does not impact the predicted ligand binding pocket and is associated with the least severe plumage depigmentation phenotype.

### Group 3: a putative regulatory change at the *EDNRB2* locus is associated with piebalding

In pigeons from Group 3, which includes both predominantly white “shield marked” breeds like the Old German Owl founder of the Hom x OGO cross and other piebald birds with less extensive white plumage, we identified 33 piebalding candidate SNPs spanning 105 kb (scaffold 507: 11145063-11249818) (Fig. 4B, Fig. 6A). We observed extended shared homozygous haplotypes in subsets of Group 3 birds, but opted not to evaluate these regions further as both small sample size and the interrelatedness of breeds in this group would make it difficult to rule out population stratification as a driver of differentiation.

**Figure 6.**
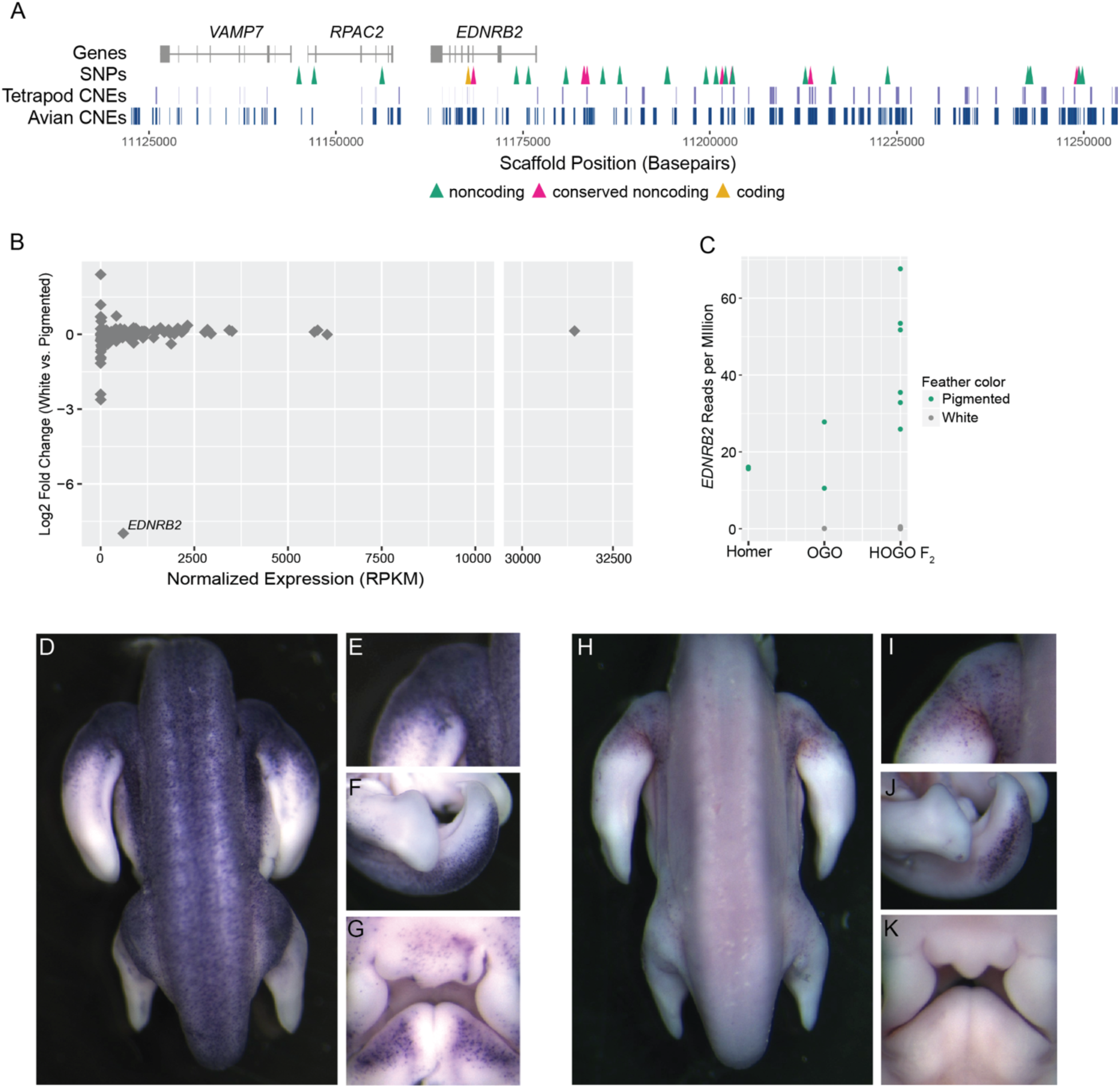
*EDNRB2* expression is altered in piebald birds. (A) Genomic context of the Group 2 piebald haplotype. Gene models for the region are shown in gray. Arrowheads denote SNPs in the following regions: yellow, coding regions; pink, non-coding regions that overlap with conserved non-coding elements; green, other noncoding regions. Conserved noncoding elements (CNEs) from tetrapod (upper) and avian-only (lower) alignments are shown in blue. (B) Differential expression (by RNA-seq) of genes within the Hom x OGO LG15 candidate region comparing white vs. pigmented regenerating feather buds from Hom x OGO F_2_ pigeons. One gene, *EDNRB2* is significantly downregulated in white feather buds. (C) Normalized *EDNRB2* read count from RNA-seq in adult Racing Homers (pigmented feather buds only), Old German Owls (OGO, pigmented and white feather buds), and Hom x OGO F_2_s (HOGO F_2_, pigmented and white feather buds). (D-K) Whole-mount *in situ* hybridization for *EDNRB2* in Racing Homer (D-G) and Classic Old Frill (H-K) embryos. Panels show dorsal body (D,H), Proximal forelimb (E,I), tail (F,J), and facial primordia (G,K).

Of the 33 piebalding candidate SNPs in Group 3, one was a coding SNP in exon 4 of *EDNRB2.* This coding mutation is synonymous and thus not likely to impact EDNRB2 function. We next examined the remaining 32 fixed noncoding SNPs to identify putative regulatory variants. We aligned genomic sequence from pigeon to orthologous regions in 15 other avian species and 6 non-avian species to identify evolutionarily conserved noncoding elements among birds and tetrapods, respectively. We defined conserved noncoding elements (CNEs) as regions with >70% sequence identity in 10 or more avian species (avian CNEs) or 3 or more non-avian tetrapods (tetrapod CNEs) [25]. We identified 9 SNPs in avian CNEs, 3 of which also overlap with tetrapod CNEs (Fig. 6A). Five of these SNPs were predicted to result in loss of one or more transcription factor binding sites [26]. One SNP in a CNE conserved in 14 of 15 avian species is predicted to result in loss of an XBP1 binding site upstream of *EDNRB2.* This is particularly relevant to piebalding as *XBP1* is linked to pigment loss as a candidate gene for vitiligo in humans, and loss of a predicted XBP1 binding site near *EDNRB2* is associated with plumage depigmentation in ducks [27,28]. Based on these findings, we hypothesized that the Group 3 haplotype includes regulatory variants that could cause plumage depigmentation by impacting the expression of *EDNRB2* or other candidate region genes.

To test this hypothesis, we examined expression of candidate genes by RNA-seq in Hom x OGO F_2_ individuals (n=6). We separately sequenced mRNA from white and pigmented regenerating feather buds from adult birds and performed differential expression analysis to quantify changes in gene expression between pigmented and non-pigmented feather buds. This F_2_ cross was founded by an Old German Owl, a representative breed from Group 3, and piebald F_2_ birds are expected to carry the Group 3 haplotype at the LG15 locus. While the fixed SNPs shared by Group 3 birds delineate a narrow candidate region spanning portions of three genes, we took a conservative approach and quantified gene expression across the broader LG15 candidate region identified by the Hom x OGO QTL analysis, which includes 112 genes. We found that only one gene within the LG15 candidate region, *EDNRB2*, was significantly differentially expressed (Fig. 6B, C). *EDNRB2* was expressed in collar cells from pigmented feather buds but was absent in collar cells from white feather buds.

We next evaluated RNA-seq from the founder Racing Homer (non-piebald, n=2) and Old German Owl (piebald, n=2) breeds. Consistent with the results from Hom x OGO F_2_ feather buds, *EDNRB2* is expressed in pigmented Racing Homer and Old German Owl feather buds, but not in white Old German Owl feather buds (Fig. 6C); only pigmented feather buds were sampled from Racing Homers as no white feathers are present in non-piebald birds. While the level of *EDNRB2* expression in pigmented regenerating feather buds varies between individuals, our RNA-seq analyses establish that *EDNRB2* expression is lost in the white feather buds of piebald birds, while other genes in the LG15 QTL candidate region are not significantly altered. This suggests that one or more of the putative regulatory mutations in Group 3 gives rise to changes in *EDNRB2* expression associated with regional plumage depigmentation.

### Group 3 piebald embryos show reduction and spatial restriction of EDNRB2 positive cells

Our RNA-seq analyses established that *EDNRB2* is differentially expressed between white and pigmented regenerating feather buds of adult birds. One of the primary roles of *EDNRB2* is mediating pigment cell migration and proliferation during embryogenesis. Therefore, we used *in situ* hybridization to examine *EDNRB2* expression in non-piebald Racing Homer embryos and piebald Classic Old Frill (Group 3) embryos across developmental stages that include pigment cell migration (Fig. 6 D-K, Supplemental Fig. S7). Classic Old Frills have a piebalding pattern characterized by white heads, bodies, and wing flight feathers, but pigmented wing shields and tails.

We found that piebald embryos show a reduction in both the number and spatial distribution of *EDNRB2-*positive cells, particularly in body regions that are unpigmented in adults. By the equivalent of chicken stage 29 of development [29], *EDNRB2*-positive cells are present throughout the dorsal body, limbs, tail, and face in non-piebald Racing Homer embryos (Fig. 6 D-G). In contrast, stage-matched Classic Old Frills have fewer *EDNRB2-*positive cells, and these cells are restricted primarily to the forelimbs and tail bud, the two regions that show pigmented plumage in adults of this breed (Fig. 6 H-J). *EDNRB2* positive cells are largely absent from the dorsal body and face (Fig. 6H,K), which are unpigmented in adults. These expression differences between breeds indicate that piebald patterns are likely established, in part, by altering pigment cell migration and proliferation early in development, both processes in which *EDNRB2* plays a key role [30–32].

We also examined embryonic *EDNRB2* expression in Hom x OGO F_2_ embryos. We found extensive variability in both the number and distribution of *EDNRB2-*positive cells, consistent with diverse plumage patterns in the adult F_2_ population used for QTL mapping (Supplemental Fig. S7). In Hom x OGO F_2_ embryos, *EDNRB2* expression is consistent with genotype at the LG15 QTL: embryos homozygous for the Racing Homer allele (N=3) look more like wild-type Racing Homer embryos, while embryos homozygous for the Old German Owl founder allele (N=3) look more like the embryos of the piebald Classic Old Frill (a close relative of the Old German Owl [33]; both breeds are part of piebalding Group 3). Heterozygous embryos (N=4) have a range of intermediate phenotypes.

### Group 4: a structural variant is associated with “baldhead” piebalding

We next evaluated piebalding candidate SNPs in Group 4, which consists of pigeon breeds with white (“bald”) heads. We identified 448 significantly differentiated SNPs on scaffold 507 spanning 619 kb, and the non-reference allele was fixed in baldhead birds at 105 of these positions. However, pigeons homozygous for all piebald-associated non-reference alleles were present in the background set, so homozygosity for these the fixed variants was not exclusive to Group 4. In addition to the haplotype enriched in baldhead birds, we noted a region where birds in this group had a large number of heterozygous SNP genotypes and increased sequencing coverage (Fig. 7A,B). Sequencing reads that align across breakpoints and targeted PCR (Supplemental Fig. S8) indicate that these birds carry a tandem duplication of a 37-kb region spanning from intron 1 of *VAMP7* to just 5’ of *EDNRB2* (Fig. 7C).

**Figure 7.**
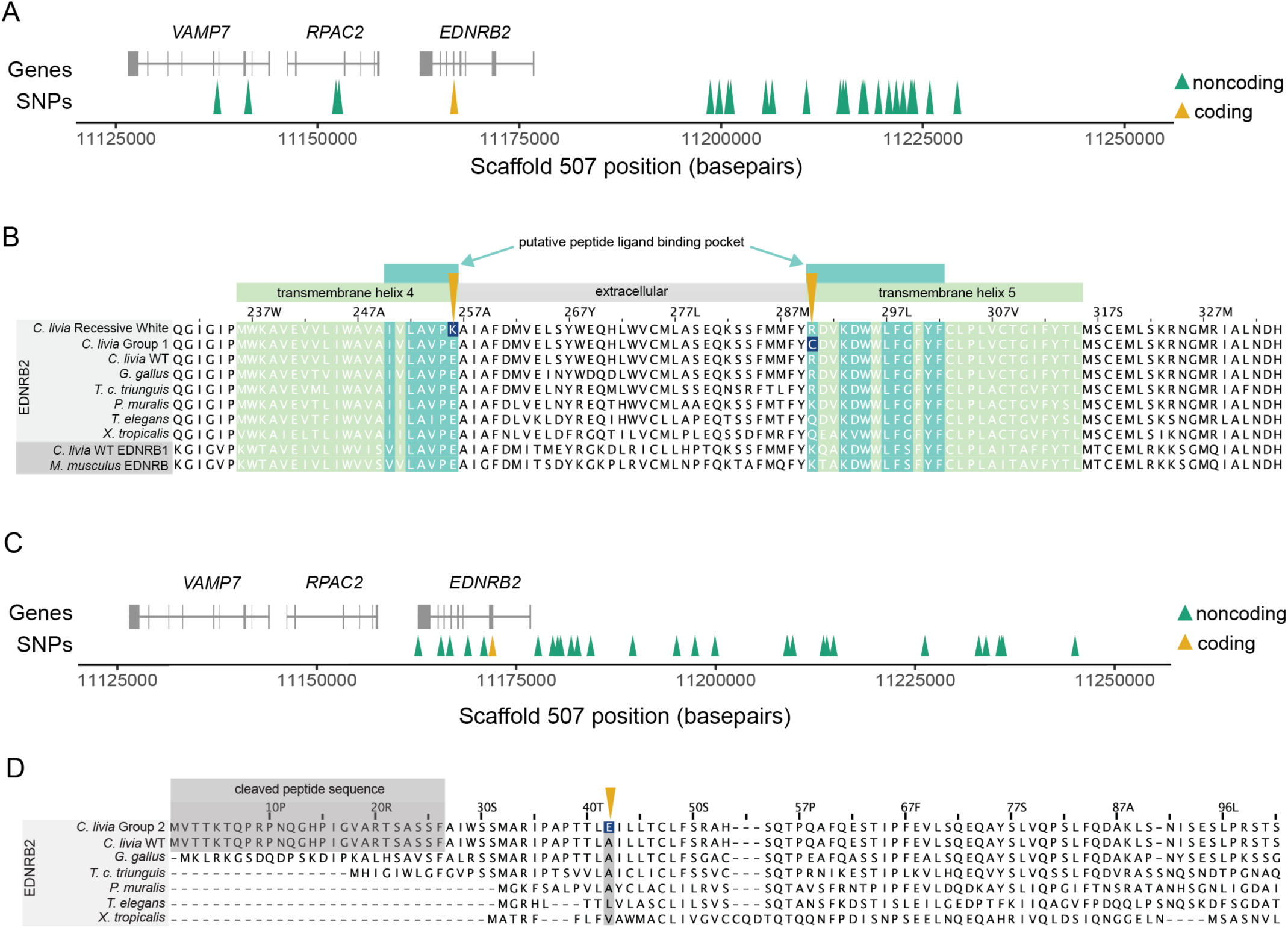
A 37-kb tandem duplication is associated with the Group 4 “baldhead” piebalding pattern. (A) Plot of genotypes within the candidate region identified by pF_ST_ for Group 4 birds. Each row shows an individual bird, and each vertical line a SNP, colored by genotype relative to the reference genome. Approximate locations of genes are illustrated at top, and regions of interest delineated by brackets, below. (B) Plot of normalized whole genome sequencing read coverage for representative birds from seven different breeds. Three baldhead breeds show an increase in coverage. * marks a coverage drop that coincides with both a LINE element and a nearby stretch of “N” bases in the reference genome and is not associated with a piebalding phenotype. (C) Schematic of the predicted structure of the baldhead-associated duplication. The duplicated region includes the 5’ portion of *VAMP7* and the entirety of both *RPAC2* and *EDNRB2.* Black triangles below indicate primer pairs used to amplify across the central breakpoint (also see Supplemental Fig. S8).

As our other SNP analyses did not assess structural variation, we screened for the 37-kb duplication in the genomes of 130 non-piebald and piebald pigeons from Groups 1-3 to see if it was associated with the Group 4 baldhead phenotype or present in non-baldhead individuals. We found two birds heterozygous and two homozygous for the duplication, but one of these homozygotes does not have a baldhead-like piebalding phenotype, suggesting that the duplication itself may not cause the characteristic Group 4 piebalding phenotype.

We did not identify any fixed coding or noncoding piebald-specific candidate SNPs in Group 4 birds. However, the presence of the tandem duplication can hinder accurate variant calling from whole genome sequencing, and variants called as heterozygous may in fact be copy-specific SNPs. Therefore, we turned to RNA-seq data from regenerating feather buds to evaluate expression of genes within the duplicated region, which could reveal regulatory variation. We first examined RNA-seq data from regenerating feather bud collar cells for two representative breeds from Group 4 that carry the duplication, the Old Dutch Capuchine and the Komorner Tumbler. Surprisingly, we found that, unlike Group 3 pigeons, *EDNRB2* is expressed in both white and pigmented regenerating feathers. However, we noted that expression in white regenerating feather buds appeared to be allele specific (Supplemental Fig. S8). Due to the duplication in this region and the high level of homozygosity at SNPs outside the duplication, we hypothesized that this expression pattern represented differential regulation of the two copies of *EDNRB2* on the same chromosome. By examining variants in reference-mapped RNA-seq data, we identified copy-specific polymorphisms that differentiate the copy of *EDNRB2* expressed in both white and pigmented feathers from the copy expressed only in pigmented feathers.

Feather buds expressing only one copy of *EDNRB2* lack pigment, suggesting that this copy is nonfunctional. This copy has four coding SNPs: three are synonymous substitutions in the white-expressed copy, and the fourth results in a proline to leucine substitution (P434L) that is not predicted to be damaging. Based on these results, coding mutations are unlikely to result in loss of function of the white-expressed *EDNRB2* copy. We next generated *de novo* transcriptome assemblies to look for potential alternative isoforms or fusion transcripts that may lack wild-type function, and identified an *RPAC2/EDNRB2* fusion transcript assembled in Old Dutch Capuchine and Komorner Tumbler *de novo* transcriptomes (baldhead Group 4 breeds), but not in Racing Homer (non-piebald) or Old German Owl (Group 3) *de novo* assemblies. RT-PCR and sequencing confirmed this fusion transcript in white feather buds from Old Dutch Capuchines and Komorner Tumblers (Supplemental Fig. S8). The predicted fusion transcript from *de novo* assemblies does not include exon 8 of *EDNRB2* and shows intronic read-through of intron 7. However, we also were able to amplify *EDNRB2* transcripts that are properly spliced across this region and include the entire coding portion of exon 8 in white feather buds from Old Dutch Capuchine and Komorner Tumblers, suggesting the presence of multiple white-expressed *EDNRB2* transcripts (Supplemental Fig. S8). While we are unable to pinpoint the precise structure of all transcripts or their potential functionality, feather buds expressing only one copy of *EDNRB2* lack pigment, indicating that these variant or fusion transcripts lack wild-type function. Additionally, the restricted expression of the wild-type non-fusion *EDNRB2* transcript in only pigmented feather buds suggests there are unidentified regulatory changes affecting the functional copy.

## DISCUSSION

### Coding and non-coding mutations in *EDNRB2* are associated with piebalding

Piebalding is a genetically and phenotypically complex suite of depigmentation phenotypes that occurs across many breeds of domestic pigeons. Our QTL mapping studies identify at least four loci that contribute to piebald patterning. Three of these are specific to different crosses, while one locus on LG15 is a major QTL for piebalding in all crosses, indicating that complex piebald patterns are determined by a combination of breed-specific and shared genetic loci (Fig. 2, Supplemental Figures S2-S5, Table 1). Among different crosses, the piebald body regions associated with the LG15 QTL are not always the same. For example, LG15 is associated with white plumage on the dorsal and lateral neck in the Hom x OGO cross, but not in the Arc x Cap or Hom x Cap crosses.

Within the shared candidate region, all piebald birds do not share a single haplotype. Instead, we identified multiple haplotypes enriched in, or specific to, subgroups of piebald birds. While the piebald-associated variants differ among groups, all likely alter the expression or function of a single candidate gene, *EDNRB2*. We identified a mix of coding variants, a putative cis-regulatory variant, and a structural variant at this locus, indicating that multiple classes of genetic change at a single locus likely contribute to diverse piebalding patterns in pigeons.

While some of the diversity of piebalding patterns is likely driven by a series of *EDNRB2* alleles, many pigeons sharing the same *EDNRB2* mutation show differences in pigment pattern. For example, pigeons homozygous for the R290C mutation in *EDNRB2* typically have pigmented heads and tails and white bodies, but pigment on the wing shield varies. Likewise, Group 3 birds carrying the same putative regulatory SNP show variation in tail pigmentation. This is consistent with the presence of additional piebalding-associated QTLs in three of our crosses. We propose that the extensive diversity of piebald patterns across domestic pigeons is controlled by the combination of *EDNRB2* allelic heterogeneity and interacting breed-specific modifiers, whose molecular identities remain unknown. Future work will further evaluate the contributions of cross-specific modifiers from QTLs on LG13 (Arc x Cap cross), LG14 (Hom x OGO), and LG1 (Pom x Scan).

### Multiple biological mechanisms could give rise to regionalized depigmentation

The establishment of epidermal pigment patterning is a complex process. Epidermal pigment cells are neural crest-derived, and multiple mechanisms can contribute to pigment loss, including defects in cell migration, proliferation, differentiation, function, and survival [4,34,35]. Upregulation of *EDNRB2* is required for pigment cell precursors to enter the dorsolateral migration pathway, mediated in part by a chemotactic response to the *EDN3* ligand [30,36]. *EDN3* also promotes melanoblast proliferation, drives ectodermal invasion, and promotes differentiation, indicating that *EDNRB2* is critical for all these processes [32,37,38]. Thus, any changes in *EDNRB2* expression or function could result in global or local plumage depigmentation.

The *EDNRB2* haplotypes we identified are associated with an array of depigmentation phenotypes. The most severe phenotype, recessive white, is associated with a protein coding change at a highly conserved residue within the putative ligand binding pocket of EDNRB2. Pigeons homozygous for this mutation lack plumage pigment and have bull eyes. Pigeon breeders have long noted an association between white plumage and bull eyes, and we previously mapped bull eye color to the *EDNRB2* locus [18]. Bull eyes appear dark due to a loss of two types of pigment in the iris stroma, allowing the dark retinal pigment epithelium at the back of the eye to show through [39]. Epidermal and iris stromal pigment are neural crest-derived, while the retinal pigment epithelium is not. Total loss of epidermal and iris stromal pigment could arise from failure of neural crest cells to enter the dorsolateral migration pathway or from widespread failure of melanoblast survival and differentiation. Future work examining expression of *EDNRB2* and other melanoblast markers in recessive white embryos could distinguish between these possibilities.

While the recessive white mutation ablates all epidermal pigmentation, a major question that arises from our study of piebald pigeons is how regionally restricted changes in depigmentation are established. For example, we found coding mutations associated with regional effects on plumage pigment, such as the *EDNRB2* R290C substitution in Group 1 birds. This could arise from a combination of changes in ligand-receptor binding affinity and non-uniform expression of ligand throughout the developing embryo, altering local interaction dynamics between piebald-associated EDNRB2 variants and the EDN3 ligand. Such changes could impact many facets of pigment patterning, including migration, proliferation, invasion, or differentiation of pigment cells. Future assessment of *EDN3* expression and the quantity and distribution of *EDNRB2* positive cells in birds harboring the R290C mutation could determine if one or all of these processes are affected.

We also identified haplotypes at the *EDNRB2* locus that are not expected to impact protein function. Instead, Group 3 piebald pigeons share a putative regulatory mutation within an evolutionarily conserved noncoding element 5’ of *EDNRB2,* which could affect pigment patterning by altering the timing, location, or level of *EDNRB2* expression. Changes in the timing or level of *EDNRB2* expression caused by regulatory mutations could impact both number and spatial distribution of migrating melanoblasts or mature melanocytes. In mice, *KIT*, not *EDNRB,* regulates melanoblast migration and differentiation. Experiments examining the effects of c-KIT inhibition in developing mouse embryos shows differentiation is spatially nonuniform, and disrupting KIT at different times leads to spatially distinct depigmentation patterns [40]. Temporally specific changes in *EDNRB2* expression could produce similar outcomes in some piebald pigeons. As *EDNRB2* plays key roles in migration, proliferation, and differentiation, other mechanisms could also give rise to breed specific patterns. For example, the specific *EDNRB2* expression pattern observed in Classic Old Frill embryos might arise from decreased proliferation coupled with a chemotactic response to long-range cues, such as EDN3 or Fibroblast Growth Factors, resulting in a reduced number of pigment cells accumulating in specific body regions [30,41].

### Endothelin B signaling mutations are associated with depigmentation in other vertebrates

Endothelin signaling is linked to changes in pigment patterning in numerous vertebrate species. Among birds, for example, tyrosinase-independent mottling in chickens, “spot” plumage in ducks, and “panda” plumage in quail are all associated with identical, convergently evolved substitutions in the sixth transmembrane domain of EDNRB2 [42–44]. Another coding mutation, in the extracellular loop between the fourth and fifth transmembrane domains, is associated with predominantly white plumage and dark eyes in Minohiki chickens [44]. This mutation does not result in fully white feathers (some dark spots persist), but like the recessive white mutation in pigeons, it impacts both plumage and iris color.

Recurrent mutations in the endothelin signaling pathway associated with pigmentation changes are not limited to birds. Zebrafish do not have a syntenic *EDNRB2* ortholog, but *EDNRBA* is critical for pigment cell migration and maturation [45]: an allelic series at the “rose” locus (*EDNRBA)* produces a range of pigmentation phenotypes, and includes missense, nonsense, and uncharacterized mutations [46]. Given the repeated involvement of *EDNRB2* in piebalding in genetically tractable domestic species, this gene should also be considered a viable candidate for pigment pattern variation among wild species. Plumage patterns featuring depigmented patches are common in myriad wild bird species, including magpies, from which the term “pied” is derived.

### Pleiotropy may constrain or permit the generation of allelic series

Endothelin pathway mutations are well documented in mammals, and they are often associated with pleiotropic effects. The endothelin signaling pathway evolved through multiple rounds of gene duplication, and expression of different combinations of endothelin receptors and ligands characterize unique cell populations [59]. In therian mammals, one endothelin B receptor copy was lost, and a single endothelin B receptor, *EDNRB,* remains. In humans, mice, and horses, *EDNRB* mutations are associated with changes in pigmentation but are coupled with deleterious effects on enteric nervous system development [9,10,14,60]. The allelic series associated with piebalding in pigeons likely would not have arisen without subfunctionalization of *EDNRB1* and *EDNRB2* that uncouples pigment patterning from enteric nervous system development. More broadly, the retention of *EDNRB2* in birds may have been crucial for the evolution of the diverse, heritable piebalding patterns of domestic pigeons, ducks, chickens, and quail.

Like *EDNRB2,* mutations in *MC1R* and *TYRP1* that alter pigmentation are also common across species [48,52,54,61–66]. In both cases, there are limited pleiotropic effects, though some *TYRP1* mutations in mice are associated with progressive hearing loss related to melanocyte degeneration and *MC1R* mutations are associated with altered immune response and analgesia [67,68]. Some *TYRP1* mutations may even confer a selective advantage, as seen in the “cinnamon” morph of American black bears [62].

The subfunctionalization of *EDNRB2* in non-mammalian vertebrates – and the developmentally-restricted functions of other pigment genes like *TYRP1* – may permit pigment diversification more readily than other traits. Interestingly, analysis of teleost fish genomes suggests that pigment genes may have been preferentially retained following genome duplication, and that this may have been a driving force behind the diversification of teleost pigment types and patterns [69]. Whole genome duplications provide the opportunity for subfunctionalization or neofunctionlization of duplicated genes while limiting the negative impacts of pleiotropy. This suggests that lack of pleiotropy may be a driving force in pigment diversification across both domestic and wild species [2,70,71].

### Multiple mechanisms may drive repeated mutation at the *EDNRB2* locus

*EDNRB2* may be a repeated mutational target in avians for a variety of reasons. Not only are pigment patterns highly visible and readily subject to selection, but subfunctionalization of the two *EDNRB* paralogs in avians also limits the pleiotropic effects of *EDNRB2* mutations. This allows for mutations to persist in populations and may provide a larger effective mutational target: coding changes, *cis-*regulatory changes, and structural variants are all potentially survivable or advantageous. In genes with multiple critical roles in development, *cis-*regulatory variants are common as altering modular regulatory elements can circumvent deleterious pleiotropic effects [70].

In some cases, the sequence content of genes or their regulatory regions can also mediate repeated mutation [72,73]. A preliminary evaluation of the *EDNRB2* region in the Cliv_2.1 assembly reveals an enrichment of both annotated transposable elements and assembly gaps (typically indicative of repetitive sequence) that may make this region more prone to mutation (Supplemental Fig. S9). Future work examining *EDNRB2* in wild avian species could determine whether convergent mutations are found across species and whether the same suite of pigment genes that drive intraspecific variation in domestic pigeons, chickens, quail, and ducks also impact interspecific variation in wild species.

## MATERIALS AND METHODS

### Animal husbandry and phenotyping of F_2_ offspring

Pigeons were housed in accordance with protocols approved by the University of Utah Institutional Animal Care and Use Committee (protocols 10-05007, 13-04012, 19-02011, and 22-03002).

Four intercrosses were used in these studies. An intercross between a male Archangel and a female Old Dutch Capuchin generated 98 F_2_ offspring [18]. An intercross between a male Racing Homer and a female Old Dutch Capuchine generated 82 offspring. An intercross between a male Racing Homer and a female old German Owl generated 171 F_2_ offspring [74]. An intercross between a male Pomeranian Pouter and two female Scandaroons generated 131 F_2_ offspring [75].

### Whole Genome Resequencing

DNA was extracted from blood samples collected with breeders’ written permission at the annual Utah Premier Pigeon Show or from our lab colony using the Qiagen DNEasy Blood and Tissue Kit (Qiagen, Valencia, CA). Samples were treated with RNAse during extraction. Isolated DNA was submitted to the University of Utah High Throughput Genomics Shared Resource for library preparation using the Illumina Tru-Seq PCR-Free library kit. The resulting libraries were sequenced on either the Illumina HiSeq or Illumina NovaSeq platforms. Raw sequence data for thirteen previously unpublished samples are available in the NCBI Sequence Read Archive under BioProject accession PRJNA986561. These data sets were combined with previously published data sets (BioProject accessions PRJNA513877, PRJNA428271, and PRJNA167554, PRJNA680754) for variant calling.

### Genomic Analyses

Variant calling was performed with FastQForward, which wraps the BWA short read aligner and Sentieon (sentieon.com) variant calling tools to generate aligned BAM files (fastq2bam) and variant calls in VCF format (bam2gvcf). Sentieon is a commercialized variant calling pipeline that allows users to follow GATK best practices using the Sentieon version of each tool [76]. FastQForward manages distribution of the workload to these tools on a compute cluster to allow for faster data-processing than when calling these tools directly, resulting in runtimes as low as a few minutes per sample.

A step-by-step summary of the workflow is available at support.sentieon.com/manual/DNAseq_usage/dnaseq/. Briefly, Raw sequencing reads from resequenced individuals were aligned to the Cliv_2.1 reference assembly [77] using fastq2bam, which utilizes the default settings of the BWA aligner. Reads are then de-duplicated using samblaster. Variant calling was performed on 199 resequenced individuals, including 186 previously published samples [47,49,50,74,75], using bam2gvcf with the quality filter “--min_base_qual 20”, and individual genome variant call format (gVCF) files were created. Joint variant calling was performed on all 186 individuals using the Sentieon GVCFtyper algorithm.

The subsequent variant call format (VCF) file was used for pF_ST_ analysis using the GPAT++ toolkit within the VCFLIB software library (https://github.com/vcflib). pF_ST_ uses a probabilistic approach to detect differences in allele frequencies between populations using a modified likelihood ratio test that incorporates genotype likelihood information [19,78]. For piebald vs. non-piebald pF_ST_ analysis, the genomes of 43 piebald birds from 31 breeds were compared to the genomes of 98 non-piebald birds (44 breeds and feral pigeons). Three breeds had both piebald and non-piebald representatives included in the analysis. The threshold for genome-wide significance was determined by Bonferroni correction (a threshold of 0.05 / total number of SNPs assayed).

### Plumage phenotyping

Individual birds from F_2_ crosses were quantitatively phenotyped. Following euthanasia, photos were taken of F_2_ plumage including dorsal and ventral views with wings and tail spread, and lateral views with wings folded. We divided the body into 15 different regions for phenotyping: dorsal head, right lateral head, left lateral head, dorsal neck, ventral neck, right lateral neck, left lateral neck, dorsal body, ventral body, dorsal tail, ventral tail, dorsal right wing, dorsal left wing, ventral right wing, and ventral left wing. To score each region, we imported photos into Photoshop v21.1.0×64 (Adobe, San Jose, CA) and used the magic wand tool to select only the white feathers within the body region. Following this selection, we saved two separate images: one containing the entire region (both pigmented and white feathers) with the color for the white feathers inverted (hereafter, “whole region image”), and one with the selected white feathers removed and only the pigmented feathers remaining (“pigmented region image”). For each body region, we imported these two images into ImageJ (v1.52a; [79] and converted them to greyscale, then used the threshold tool to measure the number of pixels in each image. To calculate the proportion of white feathers for each region, we subtracted the number of pixels in the pigmented region image from the number of pixels in the whole region image, then divided by the number of pixels in the whole region image.

Samples used for pF_ST_ were qualitatively scored as “piebald” or “non-piebald” from photographs, with published breed standards used as a reference for expectations. Birds with fully white plumage or unclear phenotyping photographs were excluded from pF_ST_ analysis, as were birds with white plumage only on the flight feathers, since breeders classify white flights as a separate phenotype unrelated to piebalding.

### Genotype-by-sequencing

DNA samples from founders of the crosses and their F_2_ progeny were extracted using the Qiagen DNeasy Blood and Tissue kit. Our genotype-by-sequencing (GBS) approach was adapted from a previously published protocol with minor modifications [75,80]. DNA was digested with ApeKI, and size selected for fragments in the 550-650 bp range. Domyan et al. (2016) performed an initial round of genotyping for the Pomeranian Pouter x Scandaroon cross. These libraries were sequenced using 100- or 125 bp paired-end sequencing on the Illumina HiSeq2000 platform at the University of Utah Genomics Core Facility. Genotype by sequencing for the Archangel x Capuchin founders (n=2) and F_2_ offspring (n=98), Homer x Capuchin founders (n=2) and F2 offspring (n=82), and Homer x Old German Owl founders (n=2) and F_2_ offspring (n=171), as well as supplemental sequencing for 20 additional and 17 previously low-coverage Pomeranian Pouter x Scandaroon F_2_s, was performed by the University of Minnesota Genomics Center. New GBS libraries were sequenced on a NovaSeq 1×100 SP flow cell.

### Linkage Map Construction

GBS reads were trimmed using CutAdapt [81], then mapped to the Cliv_2.1 reference genome reads using Bowtie2 [82]. Genotypes were called using Stacks2 by running “refmap.pl” [83].

We constructed genetic maps using R/qtl v1.46-2 (www.rqtl.org) [84]. Autosomal markers showing significant segregation distortion (*p* < 0.01 divided by the total number of markers genotyped, to correct for multiple testing) were eliminated. Sex-linked scaffolds were assembled and ordered separately, due to differences in segregation pattern for the Z chromosome. Z-linked scaffolds were identified by assessing sequence similarity and gene content between pigeon scaffolds and the Z chromosome of the annotated chicken genome (Ensembl Gallus_gallus-5.0).

Pairwise recombination frequencies were calculated for all autosomal and Z-linked markers. Markers with identical genotyping information were identified using the “findDupMarkers” command, and all but one marker in each set of duplicates was removed. Within individual Cliv_2.1 scaffolds, markers were filtered by genotyping rate; to retain the maximal number of scaffolds in the final map, an initial round of filtering was performed to remove markers where fewer than 50% of birds were genotyped. Large scaffolds (> 40 markers) were subsequently filtered a second time to remove markers where fewer than 66% of birds were genotyped.

Within individual scaffolds, R/Qtl functions “droponemarker” and “calc.errorlod” were used to assess genotyping error. Markers were removed if dropping the marker led to an increased LOD score, or if removing a non-terminal marker led to a decrease in length of >10 cM that was not supported by physical distance. Individual genotypes were removed if they had error LOD scores >5 (a measure of the probability of genotyping error, see [85]. Linkage groups were assembled from both autosomal markers and Z-linked markers using the parameters (max.rf 0.15, min.lod 6). Scaffolds in the same linkage group were manually ordered based on calculated recombination fractions and LOD scores. Linkage groups in the Pomeranian Pouter x Scandaroon map were numbered by marker number. Linkage groups in the Archangel x Old Dutch Capuchine, Homer x Old Dutch Capuchine, and Homer x Old German Owl maps were numbered based on scaffold content to correspond with Pomeranian Pouter x Scandaroon linkage groups.

### Quantitative Trait Locus Mapping

We performed QTL mapping using R/qtl v1.46-2 [84]. We used the *scanone* function to perform a single-QTL genome scan using Haley-Knott regression. For each phenotype, the 5% genome-wide significance threshold was calculated by running the same *scanone* with 1000 permutation replicates. For each significant QTL peak, we calculated 2-LOD support intervals using the *lodint* function. We calculated percent variance explained (PVE) using the *fitqtl* function.

### Identification of evolutionarily conserved non-coding regions

We identified syntenic regions spanning *ZNF185* to *TMHLE* in fifteen avian species (*Athene cunicularia, Anas platyrhynchos, Calypte anna, Corvus brachyrhynchos, Cuculus canorus, Coturnix japonica, Falco peregrinus, Gallus gallus, Haliaeetus leucocephalus, Hirundo rustica, Manacus vittelinus, Nipponia nippon, Pygoscelis adeliae, Struthio camelus,* and *Taeniopygia guttata*) and six non-avian tetrapods (*Alligator mississippiensis, Chrysemys picta bellii, Crocodylus porosus, Podarcis muralis, Thamnophis elegans,* and *Xenopus tropicalis*) using NCBI ortholog annotations and genome browser records. This region contains *EDNRB2* and its 5’ intergenic space, and encompasses the entire candidate region identified by pF_ST_. We pulled FASTA sequence from the syntenic region from the NCBI genome browser for each species and aligned these sequencing using mVISTA [25]. We defined a conserved non-coding element as any region at least 100 bp in length conserved with >70% sequence identity in 10 or more avian species (avian CNEs) or 3 or more non-avian tetrapods (tetrapod CNEs) that do not overlap annotated exons.

Evolutionarily conserved regions containing SNPs were identified by comparing the coordinates of significant SNPs fixed in Group 3 birds to tetrapod and avian CNE intervals. We generated FASTA files for CNE intervals containing both the reference and non-reference alleles using the bedtools *getfasta* function (v. 2.28.0) [86], and used TransFac Match 1.0 (http://gene-regulation.com/cgi-bin/pub/programs/match/bin/match.cgi) [26] to predict transcription factor binding sites separately for reference allele CNE sequence and alternate allele CNE sequence. We then compared lists of predicted transcription factor binding sites to identify predicted binding motifs that are gained or lost in Group 3 birds.

### Collection of collar cells from regenerating feather buds

Collar cells were collected as described previously [50]. Briefly, secondary covert wing feathers were plucked to stimulate regeneration and allowed to regenerate for 9 days. After nine days, regenerating feather buds were plucked and the proximal 5 mm was cut, bisected, and stored in RNA later at 4°C overnight. Feather buds were dissected and collar cells removed; cells were then stored at −80°C until RNA isolation.

### Expression analysis from RNA-seq data

RNA from the collar cells of regenerating feather buds was isolated using the Qiagen RNEasy Kit (Qiagen, Valencia CA), treated with the Zymo PCR Inhibitor Removal kit (Zymo Research, Irvine CA) to remove melanin, and submitted to the University of Utah High Throughput Genomics Shared Resource for Illumina TruSeq stranded library preparation. Libraries were sequenced on the Illumina NovaSeq platform. Data are available in NCBI Sequence Read Archive under BioProject PRJNA986561.

We mapped reads to the Cliv_2.1 reference genome using STAR [87], and counted reads in features using FeatureCounts [88]. We assessed differential expression in Hom x OGO F_2_ samples with DESeq2 [89]. Due to the small sample size for each breed-specific sample, we did not perform differential expression analysis. For genes of interest, we calculated and plotted normalized gene expression based on the number of uniquely mapped reads within the gene model and the total number of reads per sample.

### Protein conservation, structure prediction, and mutation evaluation

We obtained protein sequences for EDNRB2 orthologues across species using NCBI blastp and generated multi-species alignments using Clustal Omega [90], and then visualized using Jalview2 [91]. The boundaries of conserved domains were determined from the NCBI Conserved Domain Database for protein accession number cd15977 (https://www.ncbi.nlm.nih.gov/Structure/cdd/cddsrv.cgi?uid=320643, last access date 8/17/2022) [21,22]. The probable impact of coding mutations were evaluated using the PolyPhen2 web server (http://genetics.bwh.harvard.edu/pph2/, access date 8/17/2022) [23].

### Identification and evaluation of structural variants

We identified reads with aberrant insert sizes within the scaffold 507 candidate region using both the Samtools suite and visualization with the IGV genome browser [92,93]. In a subset of samples, we observed large pileups of read pairs with larger than normal insert sizes at two loci, within *VAMP7* and upstream of *EDNRB2*, and an apparent increase in coverage between them. We also identified numerous soft-clipped reads within *VAMP7* and directly upstream of *EDNRB2* and used NCBI BLAST to compare this soft-clipped sequence to the Cliv_2.1 reference genome [77,94]. Mapping of this soft-clipped sequence suggested a tandem duplication. We next calculated coverage across 5 kb sliding windows on scaffold 507 using the Bedtools *genomecov* function (v2.28.0) [86]. Coverage was normalized to 2x and plotted. We designed primers near the predicted duplication breakpoints (Left TATGCAGCAGCACACAAACA; Right GCAGTCACTCGTTTCCCTCT) and used PCR to amplify across the central breakpoint of the tandem duplication to confirm its presence.

### *De novo* transcriptome assembly and analysis

Transcriptomes were assembled from collar cell RNAseq data using Trinity v. 2.11.0 with the “--jaccard-clip” option [95]. Transcripts fully or partially matching *EDNRB2* were identified by NCBI Blast [94], using the *de novo* transcriptome assembly as the database. These transcripts were then aligned to the reference *EDNRB2* mRNA sequence (accession XM_021292346.1) using Clustal Omega [96]. Portions of assembled transcripts that did not align to *EDNRB2* mRNA were compared to the Cliv_2.1 reference genome and transcriptome using NCBI Blast to evaluate sequence similarity to intronic and intergenic sequence as well as other annotated genes. The predicted fusion transcript between *EDNRB2* and *RPAC2* was then validated by RT-PCR.

### RT-PCR

RNA from collar cells was isolated as for RNA-seq. cDNA was synthesized from isolated collar cell RNA using the Invitrogen SuperScript IV first strand synthesis system and Oligo dT primer (Invitrogen #18091050). We then used this cDNA as input for PCR. *EDNRB2* expression was assessed using a reverse primer in *EDNRB2* exon 8 (CCCCAGGTTATGTTGGTCAC) and a forward primer in *EDNRB2* exon 7 (CCCTGTGGCTCTCTACTTCG). The presence of an RPAC2/EDNRB2 fusion transcript was confirmed using a reverse primer in *EDNRB2* (3’ to 5’ in exon 2 of *EDNRB2;* GCCAAGCTAGCAATGAGGAC) and forward primer in *RPAC2* (3’ to 5’ in intron 1 of *RPAC2*; GGGTTGCAACATCTGCACTA).

### *In situ* hybrization

ISH probe templates were generated by PCR amplification using primers targeting a portion of pigeon *EDNRB2* (279-bp amplicon, Fwd ATGAGGAGGAGAGGGAGAGG; Rev AAGGTGAAGAGAAGGGGTGG) from a cDNA library generated from pooled feather bud samples from multiple breeds. The *EDNRB2* amplicon were cloned into pGEM-T Easy (Promega) and confirmed by Sanger sequencing. To generate sense and antisense probes, pGEM-*EDNRB2* was digested with NcoI and transcribed with SP6 RNA polymerase (sense) or digested with SalI and transcribed with T7 RNA polymerase (antisense).

Racing Homer, Classic Old Frill, and Hom x OGO F_2_ embryos used for ISH were dissected from eggs at the desired embryonic stage and fixed overnight in 4% paraformaldehyde at 4°C on a shaking table. Embryos were subsequently dehydrated into 100% MeOH and stored at −20°C. Whole-mount ISH was performed essentially as previously described following a protocol optimized for avian embryos (http://geisha.arizona.edu/geisha/protocols.jsp) [74].

### Repeat, transposable element, and N tract quantification

Intervals annotated as simple repeats or transposable elements were extracted from the Cliv_2.1 repeat annotation GFF [77]. Locations of N tracts were extracted in BED format using perl. Random windows size-matched to the candidate region across scaffold 507 were generated using the bedtools (v. 2.8.2) *random* function, and overlap of windows with repeats, TEs, and N tracts calculated using bedtools *intersect* [86].

## Data Availability

Whole genome sequencing and RNA-sequencing datasets generated for this study have been deposited to the NCBI SRA database under BioProject PRJNA986561 Previously generated whole genome sequencing data used in this study is available under BioProject accessions PRJNA680754, PRJNA513877, PRJNA428271, and PRJNA167554.

## Author Contributions

E.T.M. and M.D.S. designed the study. E.T.M. and A.M.S. analyzed whole-genome sequencing data. E.T.M. and E.F.B. analyzed GBS data and assembled linkage maps. E.T.M., R.W., B.P., and A.B. phenotyped animals and carried out QTL mapping. E.M. and R.W. collected and processed regenerating feather bud samples. E.T.M. and B.P. generated and analyzed RNA-seq data. E.T.M., R.W., and A.B. performed *in situ* hybridizations. M.D.S. supervised the study. E.T.M. and M.D.S. wrote the manuscript, with input from all authors.

## Acknowledgements and Funding Information

We thank current and former members of the Shapiro lab for assistance with sample collection and processing. We thank Anna Vickrey, Hannah van Hollebeke, and Alexa Davis for technical assistance and advice. Layne Gardner generously shared the Pomeranian Pouter and Scandaroon photographs in Fig. 1. We thank members of the Utah Pigeon Club for providing samples. We acknowledge a computer time allocation from the University of Utah Center for High Performance Computing. This work was supported by the National Institutes of Health (R35GM131787 to M.D.S.), a fellowship from the Jane Coffin Childs Memorial Fund for Medical Research to E.M., and the University of Utah Undergraduate Research Opportunities Program (fellowship support to B.P. and R.W.).

